# High-Purity Production of Endothelial Cells from Human Pluripotent Stem Cells

**DOI:** 10.1101/2025.07.05.663260

**Authors:** Koki Yoshimoto, Shiho Terada, Ken-ichiro Kamei

**Affiliations:** Institute for Integrated Cell-Material Sciences, Kyoto University, Kyoto, 606-8501, Japan; Program of Biology, Division of Science, New York University Abu Dhabi, Abu Dhabi, P.O. Box 129188, United Arab Emirates; Program of Bioengineering, Division of Engineering, New York University Abu Dhabi, Abu Dhabi, P.O. Box 129188, United Arab Emirates; Department of Biomedical Engineering, Tandon School of Engineering, New York University, New York, NY 10003, USA; Department of Biology, Faculty of Arts & Science, New York University, Brooklyn, NY 11201, USA; Department of Pharmaceutics, Wuya College of Innovation, Shenyang Pharmaceutical University, Shenyang, 110016, China; Joint International Research Laboratory of Intelligent Drug Delivery Systems, Ministry of Education, Shenyang, 110016, China

**Keywords:** Endothelial cells, human pluripotent stem cells, WNT, NOTCH, differentiation, organogenesis, RNA sequencing

## Abstract

Endothelial cells (ECs) are essential for vascular network formation and tissue homeostasis, yet the fields of tissue engineering and vascularized organoid generation still relies heavily on human umbilical vein ECs (HUVECs), which are venous, allogeneic, and difficult to mature fully. Human pluripotent stem cells (hPSCs) offer an autologous, developmentally flexible alternative, but most differentiation protocols require fluorescence-activated cell sorting, limiting scalability. Here we present a streamlined method that produces highly pure ECs directly from human embryonic stem cells (hESCs) without cell sorting. Extending Wnt activation with CHIR99021 to three days maximizes mesoderm induction, and brief Notch blockade with DAPT during specification suppresses smooth-muscle commitment. The result is over 90 % CD31^+^ CD144^+^ ECs that display classic cobblestone morphology, robust DiI-acetylated LDL uptake, and capillary-like sprouting comparable to HUVECs. Bulk RNA barcoding and sequencing segregates the hESC-derived ECs from both HUVECs and undifferentiated hESCs and uncovers an arterial-skewed transcriptome: *NOTCH1, DLL4*, and *CXCR4* are up-regulated, whereas venous markers (*EPHB4, NRP2*) are reduced. Enrichment of Notch-responsive pathways further supports an arterial-like identity. Although several adult functional genes (e.g., *vWF, NOS3*) are expressed at lower levels than in HUVECs, the protocol delivers a scalable source of developmentally relevant ECs ideal for vascularizing organoids derived from the same hPSCs and for future applications in drug screening, disease modeling, and cell-based vascular therapies.

## Introduction

Endothelial cells (ECs) facilitate the differentiation and maturation of parenchymal cells within tissues through paracrine signaling and direct interactions during the early stages of organogenesis.^1–5^ Subsequently, these cells line the entirety of the vascular system, forming a dynamic interface critical for the transport of oxygen, nutrients, and signaling molecules, as well as for immune surveillance and response, thereby ensuring tissue viability and function.^6^ Given these multifaceted and vital roles, the availability of functional human ECs is a cornerstone for advancing engineered human tissues, such as organoids that recapitulate *in vivo*-like complexities, and for developing novel strategies in regenerative medicine and drug discovery.^7^

Human pluripotent stem cells (hPSCs), including human embryonic stem cells (hESCs)^8^ and human induced pluripotent stem cells (hiPSCs)^9^, have emerged as a highly promising and scalable source for generating a wide array of cell types, including those needed for complex organoid assembly.^7^ For vascularization strategies in engineered tissues, human umbilical vein endothelial cells (HUVECs) have historically been a common choice due to their relative ease of isolation and commercial availability. However, the utility of HUVECs in sophisticated tissue engineering and regenerative applications is increasingly recognized as limited. HUVECs often exhibit a restricted capacity to fully support the differentiation and maturation of co-cultured parenchymal cells compared to primary microvascular or organ-specific ECs.^10,11^ Furthermore, their inherent venous identity makes them suboptimal for constructing arterialized or organ-specific vascular networks, which are critical for recapitulating physiological tissue function.^12,13^ However, HUVECs exhibit a limited ability to promote differentiation and maturation compared to primary human endothelial cells.^14^ most critically, the allogeneic nature of HUVECs presents significant immunological hurdles for their use in artificial tissues intended for therapeutic transplantation, potentially leading to rejection and limiting their clinical translatability.^15^

To address these limitations, ECs differentiated from patient-specific or HLA-matched hPSCs are gaining prominence as a superior alternative for organoid vascularization and regenerative therapies.^16,17^ hPSC-ECs offer the potential for autologous or immunologically compatible cell sources, tunable differentiation towards specific endothelial subtypes (e.g., arterial, venous, lymphatic, or organ-specific), and robust scalability.^4,5,18,19^ However, a persistent challenge in the field has been the inefficiency and heterogeneity of many current EC differentiation protocols. Many established methods yield EC populations (often identified by CD31/PECAM1 and CD144/VE-Cadherin co-expression) with purities often below 80%, necessitating subsequent enrichment steps such as fluorescence-activated cell sorting (FACS) or magnetic-activated cell sorting (MACS).^18,20^ While effective, these purification procedures add considerable time, labor, and expense to the cell production process, and can potentially impact cell viability and function, thereby hindering the scalability required for large-scale tissue engineering, drug screening platforms, and clinical applications.^21^ The establishment of a high-efficiency differentiation protocol that enables direct, sort-free induction of ECs from hPSCs is therefore of paramount importance, considering factors, such as cost, time and scalability.

Early protocols explored the direct differentiation of hPSCs into ECs by activating key signaling pathways such as MEK/ERK and BMP4, achieving modest yields of CD31^+^CD34^+^ cells (e.g., ∼15%)^22^. Subsequent research underscored the pivotal role of precise temporal modulation of WNT signaling during mesoderm specification, revealing that transient WNT activation followed by its inhibition can significantly steer mesodermal progenitors towards an endothelial fate^18,23^. Furthermore, the interplay between VEGF signaling, crucial for endothelial proliferation and survival, and Notch signaling has been identified as a key regulatory axis. Specifically, Notch inhibition, often in conjunction with VEGF stimulation, has been reported to promote the specification and expansion of endothelial progenitors from hPSCs^20^. Despite these advancements, achieving direct differentiation efficiencies consistently exceeding 80-90% without sorting has remained elusive for many protocols, often due to incomplete lineage commitment or the emergence of off-target cell types^24–26^. Recent efforts continue to refine these strategies, focusing on optimized growth factor combinations, small molecule modulators, and defined culture conditions to improve purity and yield^21,27,28^. However, the need for robust, sort-free protocols that are easily adaptable and scalable remains paramount for widespread application.

Here we report a significantly refined and highly efficient protocol for the directed differentiation of hPSCs into ECs. By systematically optimizing the duration and timing of mesoderm induction using the selective GSK-3 inhibitor CHIR99021, and strategically incorporating the Notch inhibitor DAPT during endothelial specification, our method consistently yields EC populations exceeding 90% purity for CD31 and CD144 co-expression, directly from hPSCs without any requirement for cell sorting. We demonstrate that these hPSC-derived ECs exhibit characteristic endothelial morphology, robust sprouting angiogenesis *in vitro*, comparable to HUVECs, and possess a transcriptomic signature indicative of an arterial endothelial phenotype, characterized by enriched expression of arterial markers. This optimized, sort-free approach offers a scalable, cost-effective, and time-efficient means to generate large quantities of high-purity hPSC-ECs, providing a valuable and versatile cell source for constructing vascularized organoids, advancing regenerative medicine strategies, and developing more predictive drug discovery platforms.

## Materials and methods

### Human Embryonic Stem Cell (hESC) Culture and Maintenance

Human embryonic stem cells (hESCs), line H9 (WA09; RRID: CVCL_9773), were obtained from WiCell Research Institute (Madison, WI, USA). All hESC experiments were conducted in accordance with the guidelines of the Ethics Committee of Kyoto University (Approval No. ES3-9).

For routine culture, hESC-certified Matrigel (Corning, Inc., Corning, NY, USA) was diluted 1:75 (v/v) with DMEM/F12 medium (Merck KGaA, Darmstadt, Germany) and used to coat culture dishes. The Matrigel solution was incubated in the dishes for 24 h at 4°C. Prior to cell seeding, excess Matrigel was aspirated, and the coated surface was washed with fresh DMEM/F12 medium. hESCs were maintained in mTeSR1 medium (Stem Cell Technologies, Vancouver, Canada) supplemented with 1% (v/v) penicillin/streptomycin (Fujifilm Wako, Osaka, Japan). The medium was changed daily.

For passaging, cells were first washed with D-PBS (devoid of calcium and magnesium; Thermo Fisher Scientific, Waltham, MA, USA). Dissociation was achieved by incubating the cells with TrypLE Express (Thermo Fisher Scientific) for 3 min at 37°C. The detached cells were harvested and pelleted by centrifugation at 200 ×*g* for 3 min. Cell pellets were subsequently resuspended in mTeSR1 medium. To enhance cell survival post-dissociation, mTeSR1 medium supplemented with 10 μM Y27632 ROCK inhibitor (Fujifilm Wako) was used for the first 24 h after passaging. Thereafter, cells were cultured in mTeSR1 medium without ROCK inhibitor, with daily medium replacement. Cells were maintained for a maximum of 10 passages.

### Differentiation of hESCs towards Endothelial Cells

To initiate endothelial differentiation, cultured hESCs were washed with D-PBS and detached using TrypLE Express for 3 min at 37°C. The reaction was neutralized by the addition of basal medium, and the cell suspension was transferred to a 15-mL conical tube. Cells were pelleted by centrifugation at 200 ×*g* for 3 min, and the supernatant was discarded. The cell pellet was resuspended in mTeSR1 medium supplemented with 10 μM Y27632 and 1% (v/v) penicillin/streptomycin. Cells were seeded at a density of 5.3 × 10^4^ cells cm^−2^ onto Matrigel-coated wells of Nunc Cell-Culture Treated Multidishes (Thermo Fisher Scientific). The cells were cultured in a humidified incubator at 37°C with 5% CO_2_ for 24 h.

Subsequently, the medium was replaced daily for 3 days with differentiation medium 1, consisting of DMEM/F12 (Merck KGaA) supplemented with 25 ng/mL human recombinant BMP4 (R&D Systems, Minneapolis, MN, USA), 8–16 μM CHIR99021 (concentration adjusted based on cell confluency and morphology; ReproCELL, Kanagawa, Japan), 50 ng mL^−1^ human recombinant bFGF (Fujifilm Wako), 1% (v/v) B27 supplement (Thermo Fisher Scientific), 1% (v/v) GlutaMax Supplement (Thermo Fisher Scientific), and 1% (v/v) penicillin/streptomycin.

Following this, cells were cultured for an additional 3 days with daily medium changes using differentiation medium 2, composed of StemPro-34 SFM (Thermo Fisher Scientific) supplemented with 100 ng mL^−1^ human recombinant VEGF-A_165_ (Fujifilm Wako), 10 μM DAPT (AdipoGen Life Sciences, Inc., San Diego, CA, USA), 2 μM Forskolin (Tokyo Chemical Industry Co., Ltd., Tokyo, Japan), 1% (v/v) GlutaMax Supplement, and 1% (v/v) penicillin/streptomycin.

Finally, cells were matured for 5 days in EGM Endothelial Cell Growth Medium BulletKit (Lonza, Basel, Switzerland) supplemented with 100 ng mL^−1^ VEGF-A_165_ and 1% (v/v) penicillin/streptomycin, with daily medium changes. If excessive cell proliferation was observed on day 10 of differentiation (corresponding to day 4 of EGM treatment), the medium change was omitted for that day.

### Culture of Human Umbilical Vein Endothelial Cells (HUVECs)

HUVECs (KAC Co., Ltd., Kyoto, Japan) were cultured in EGM Endothelial Cell Growth Medium BulletKit (Lonza) supplemented with 100 ng/mL VEGF-A_165_ and 1% (v/v) penicillin/streptomycin. For passaging, HUVECs were washed with D-PBS (devoid of calcium and magnesium) and dissociated using 0.02% EDTA / 0.1% trypsin solution (Kohjin-bio, Saitama, Japan) for 3 minutes at 37°C. Detachment was neutralized by the addition of EGM medium containing 1 mg mL^−1^ trypsin inhibitor from soybean (Fujifilm Wako) in PBS supplemented with 1% (v/v) penicillin/streptomycin. The cells were pelleted by centrifugation at 200 ×*g* for 3 min, resuspended, and seeded in fresh EGM medium.

### Fluorescence-Activated Cell Sorting (FACS)

Cells intended for FACS analysis were rinsed twice with PBS and harvested using 0.1% trypsin/EDTA, followed by neutralization with 1 mg mL^−1^ trypsin inhibitor. After cell counting, cells were diluted to a final concentration of 1 × 10^7^ cells mL^−1^ in staining buffer (Fetal Bovine Serum (FBS); BD Pharmingen, Franklin Lakes, NJ, USA). For antibody staining, 2.5 μL of fluorescence-labeled antibody (specific antibodies listed in Supplementary Table S1) was added to 50 μL of the cell suspension, and the mixture was incubated on ice (approximately 4°C) for 30-60 min. Corresponding isotype controls were used at the same concentration as the primary antibodies for negative controls. After incubation, excess antibodies were removed by washing the cells with staining buffer, followed by centrifugation at 300 ×*g* for 3 min. The stained cell suspensions were analyzed and sorted using a FACSAria II SORP cell sorter (BD Biosciences, Franklin Lakes, NJ, USA). Data were analyzed using FlowJo software (v9; FlowJo, LLC, Ashland, OR, USA).

### Immunocytochemistry

Cells cultured on appropriate surfaces were fixed with 4% paraformaldehyde (PFA) in D-PBS (devoid of calcium and magnesium) for 20 min at 25°C. Following fixation, cells were permeabilized with 0.1% (v/v) Triton X-100 in D-PBS for 5 min at 25°C. Non-specific binding sites were blocked by incubating the cells in blocking buffer [D-PBS containing 5% (v/v) normal goat serum (Maravai Life Sciences, San Diego, CA, USA), 5% (v/v) normal donkey serum (Jackson ImmunoResearch, West Grove, PA, USA), 3% (w/v) bovine serum albumin (BSA), essentially globulin-free (Merck KGaA), and 0.1% (v/v) Tween-20 (Nacalai Tesque, Kyoto, Japan)] at 4°C for 16 h. Cells were then incubated with primary antibodies, diluted in blocking buffer as detailed in Supplementary Table S3, at 4°C for 16 h. Subsequently, cells were washed and incubated with appropriate secondary antibodies, diluted as described in Supplementary Table S3, at 37°C for 60 min. Nuclei were counterstained with DAPI (Fujifilm Wako) at 25°C for 30 min.

### Bulk RNA barcoding and sequencing

Total RNA was purified using the RNeasy Micro Kit (Qiagen, Hilden, Germany), and stored at −80 °C until library preparation. Bulk RNA barcoding and sequencing (BRB-seq)^29^ was libraries were generated with the following modifications. First-strand cDNA was synthesized with an oligo-dT primer, and double-stranded cDNA was produced using the Second Strand Synthesis Module (New England Biolabs, #E6111). Tagmentation was carried out with an in-house MEDS-B Tn5 transposase^30,31^, followed by 10 PCR cycles using Phusion High-Fidelity DNA Polymerase (Thermo Fisher Scientific, M0530). Paired-end sequencing (Read 2 length: 81 bp) was performed on an Illumina NovaSeq 6000.

### BRB-seq Data Processing

Barcodes were extracted with UMI-tools v1.1.1 using:

“umi_tools extract -I read1.fastq --read2-in=read2.fastq ¥

--bc-pattern=NNNNNNNNNNCCCCCCCCC --read2-stdout”

Adaptor trimming, low-quality base removal, and length filtering (<20 bp) were conducted with Trim Galore v0.6.7. High-quality reads were aligned to the human reference genome (GRCh38) with HISAT2 v2.2.1, and gene-level counts were generated with featureCounts v2.0.1. Differential expression analysis was performed in DESeq2 v1.34.0 and iDEP.96^32^, applying |log_2_(fold change)| > 1 and adjusted *p* < 0.05 as significance thresholds.

### Endothelial Sprouting Assay

Dishes were coated with 100 μL of Geltrex (Thermo Fisher Scientific) and incubated at 37 °C for 30 min. Cells were seeded in EGM™ Endothelial Cell Growth Medium BulletKit and cultured for 48 h. Sprouting morphology characteristic of endothelial cells was then assessed microscopically.

### Fluorescence Imaging

Samples were imaged on a Nikon ECLIPSE Ti inverted fluorescence microscope equipped with a CFI Plan Fluor 10×/0.30 NA objective, ORCA-R2 CCD camera (Hamamatsu Photonics), Intensilight mercury lamp (Nikon), motorized XYZ stage (Ti-S-ER with encoders), and DAPI, GFP HYQ, and TRITC filter cubes (Nikon).

### Statistical Analysis

Statistical analyses were conducted in *R* v4.0.2. One-way ANOVA followed by Tukey’s post-hoc test or two-way ANOVA with Welch’s correction was used as appropriate. Differences were considered significant at *p* < 0.05.

## Results

### Notch Inhibition Enhances hPSC Differentiation into Endothelial Cells

To optimize endothelial differentiation from human pluripotent stem cells (hPSCs), we refined a published protocol (Figure 1A)^23^ by adjusting both CHIR99021, a selective GSK-3 inhibitor that promotes mesoderm induction, and DAPT, a γ-secretase/Notch inhibitor that favors endothelial specification. Using the original method, the resulting cells displayed a fibroblast-like morphology (Supplementary Figure 1A) and contained only ∼10 % CD144^+^ CD31^+^ cells (Figure 1B; Supplementary Figure 1B). In contrast, the fine-tuned protocol yielded cells with a typical cobblestone endothelial morphology (Supplementary Figure 1C) and markedly improved differentiation efficiency: at a seeding density of 1.9 × 10^4^ cells cm^−2^, 68.9 % of cells were CD144^+^ CD31^+^. Increasing the density to 5.3 × 10^4^ cells cm^−2^ further boosted efficiency, ultimately producing 90.0 % CD144^+^ CD31^+^ cells directly from hESCs without the need for cell sorting (Figure 1C; Supplementary Figure 1C).

**Figure 1:**
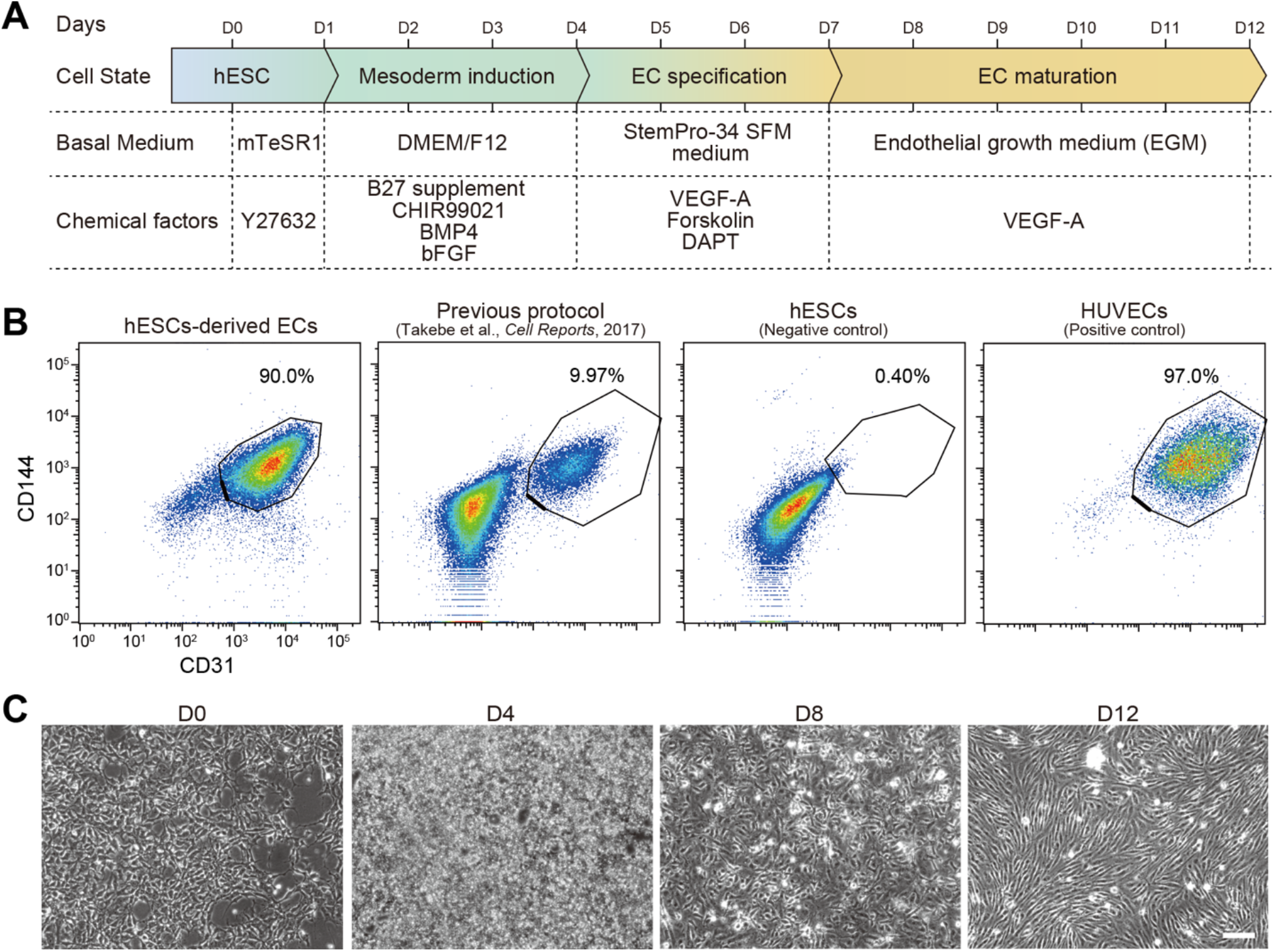
Efficient differentiation of hESCs into endothelial cells. (**A**) Schematic illustration of the protocol to generate endothelial cells (ECs) from human embryonic stem cells (hESCs): brief Wnt activation with CHIR99021 followed by Notch inhibition with DAPT. (**B**) Flow-cytometric analysis for VE-cadherin (CD144) and PECAM-1 (CD31). Shown left to right: undifferentiated hESCs (negative control), cells from the original protocol, cells from the optimized protocol, and HUVECs (positive control). Percent CD144^+^ CD31^+^ cells is noted for each group. (**C**) Phase-contrast micrographs of cells during differentiation from hESCs to ECs with the optimized method. Scale bar, 100 μm.

### hPSC-Derived Endothelial Cells Express Canonical Markers and Form Sprouts

Immunostaining confirmed robust surface expression of CD31 and CD144 in hPSC-derived endothelial cells (ECs), comparable to human umbilical vein endothelial cells (HUVECs) (Figure 2A). In a 3-D sprouting assay, the differentiated cells generated capillary-like sprouts similar to HUVECs, whereas mesenchymal control cells did not (Figure 2B).

**Figure 2:**
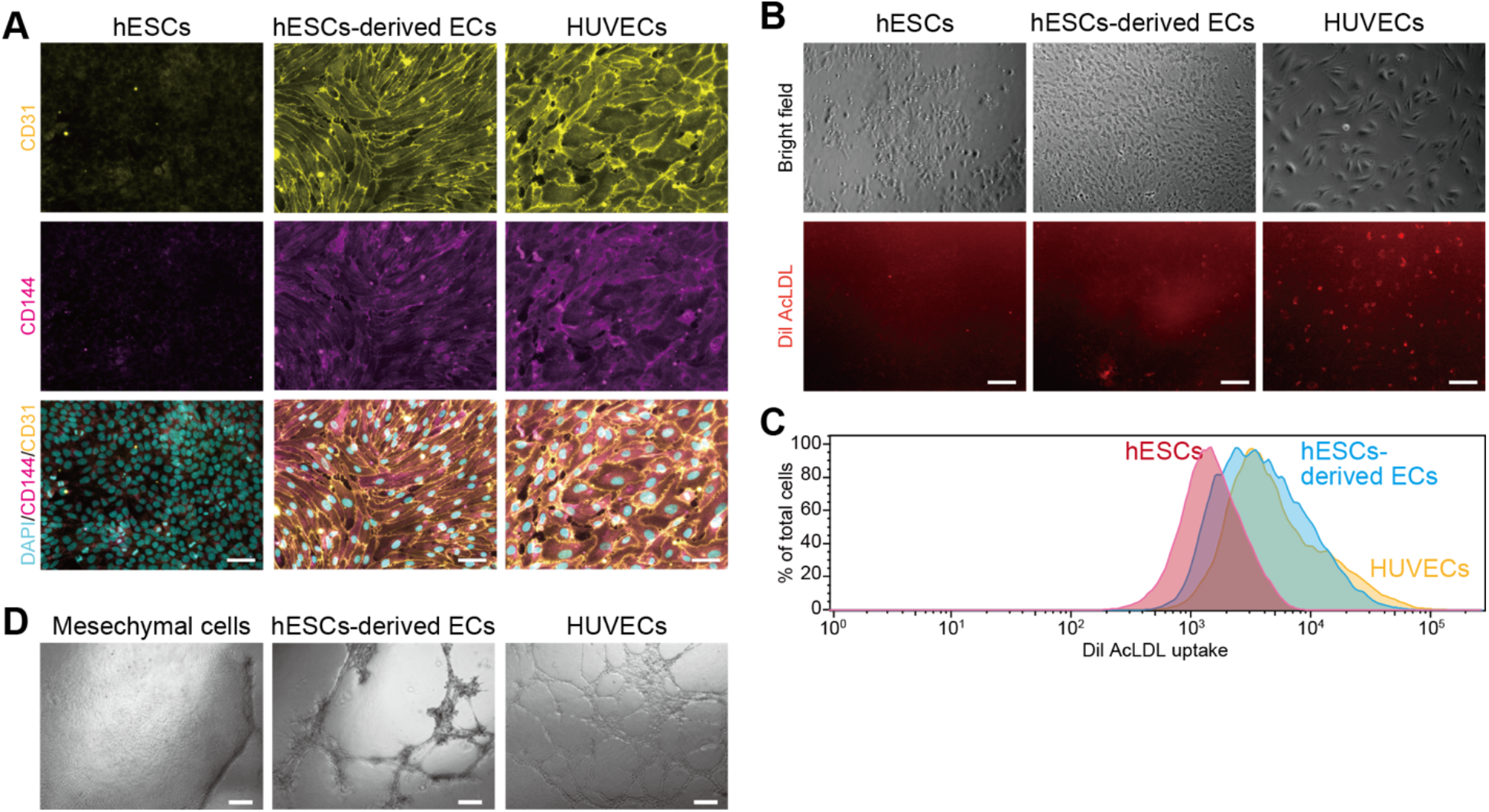
Phenotypic and functional characterization of hESC-derived endothelial cells. **(A)** Immunofluorescence staining for PECAM-1/CD31 (yellow) and VE-cadherin/CD144 (magenta); nuclei counterstained with DAPI (cyan). Undifferentiated hESCs and mesenchymal cells serve as negative controls, HUVECs as a positive control. Scale bar, 100 μm. (**B**) DiI-acetylated LDL uptake visualized by fluorescence microscopy. Scale bar, 100 μm. (**C**) Single-cell DiI-AcLDL fluorescence profiles obtained by flow cytometry confirm efficient LDL uptake by hESC-derived ECs and HUVECs. (**D**) Three-dimensional sprouting assay showing capillary-like projections formed by hESC-derived ECs and HUVECs, but not by mesenchymal controls. Scale bar, 100 μm.

### BRB-seq Reveals Distinct Transcriptional Profiles in hPSC-Derived ECs and HUVECs

Bulk RNA barcoding and sequencing (BRB-seq) followed by principal-component analysis cleanly separated hPSC-derived ECs, undifferentiated hESCs, and HUVECs into three clusters (Figure 3A). PC1 (64 % variance) reflected the pluripotent-to-endothelial transition, whereas PC2 (24 % variance) distinguished hPSC-derived ECs from HUVECs.

**Figure 3:**
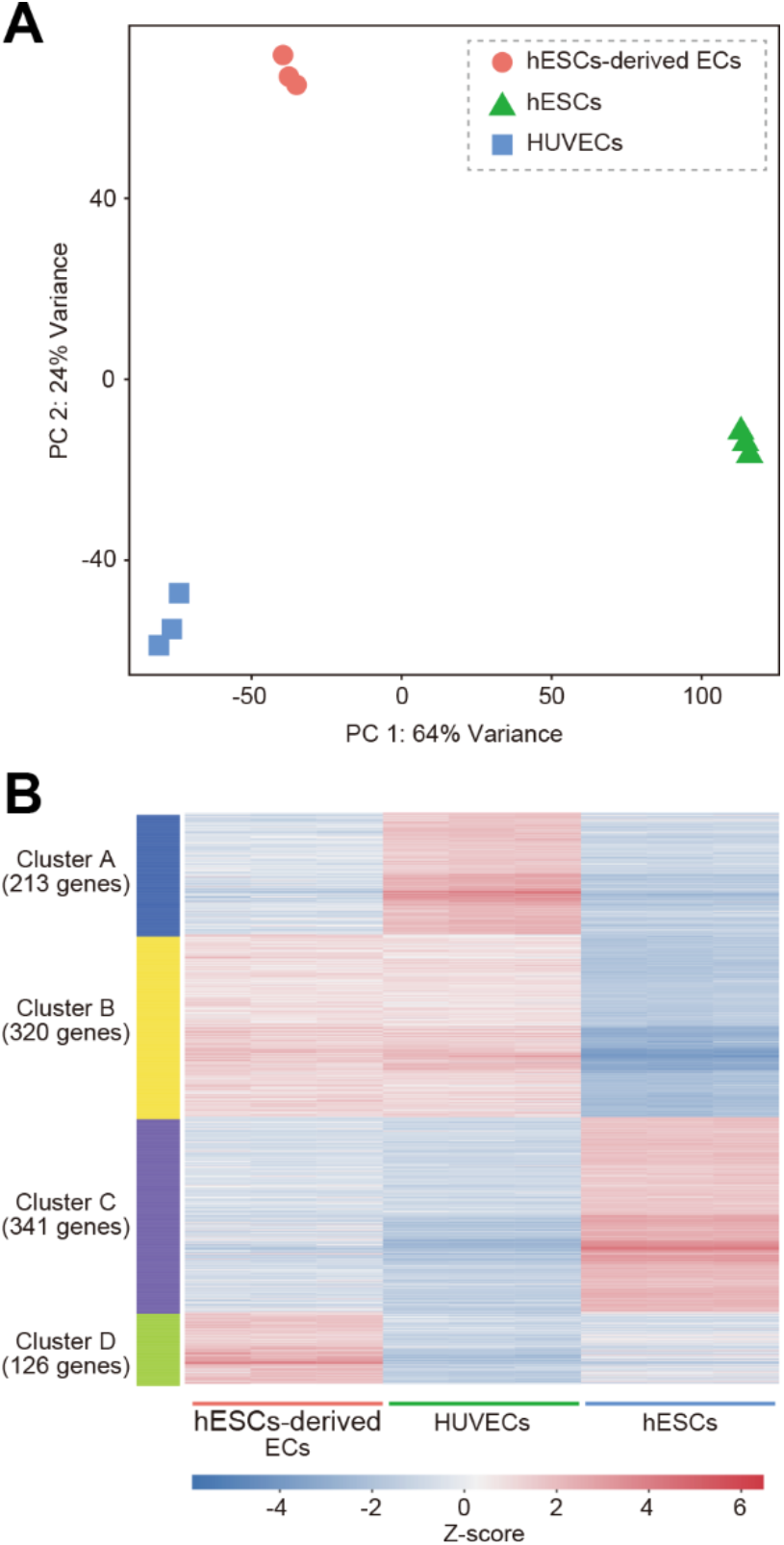
Transcriptional comparison of hESC-derived endothelial cells, undifferentiated hESCs, and HUVECs (BRB-seq). (**A**) Principal-component analysis of BRB-seq profiles cleanly separates hESC-derived ECs (pink), hESCs (green), and HUVECs (blue) into three distinct clusters. (**B**) Heatmap of the 1000 most variable genes, grouped by *k*-means clustering (*k* = 4). Each column represents a sample; each row, a gene.

K-means clustering of the 1,000 most variable genes (k = 4) identified four gene sets with distinct pathway enrichments (Figure 3B; Table 1), such as Cluster A (213 genes), enriched in HUVECs and highlighted adhesion/migration pathways (e.g., *CLDN11, COL8A1*, and *LYVE1*), Cluster B (320 genes), shared by both endothelial populations and contained classic EC genes (*CD31*, and *CDH5*) and “blood-vessel development” pathways, Cluster C (341 genes) marked pluripotent hESCs (*SOX2*, and *POU5F1*) with “cell-differentiation” signatures, and Cluster D (126 genes) specific to hPSC-derived ECs (*COL1A1*, and *SOX6*) and enriched for morphogenesis/developmental processes.

Thus, hPSC-derived ECs and HUVECs share a core endothelial program yet maintain distinct transcriptional identities.

### Arterial Gene Programs Are Up-Regulated in hPSC-Derived ECs

Differential-expression analysis between hPSC-derived ECs and HUVECs revealed higher expression of KDR and T-cadherin in the former (Figure 4A,B). Enrichment analysis of up-regulated genes pointed to “blood-vessel development” and “tube morphogenesis” pathways, featuring arterial markers such as *NOTCH1, CXCR4*, and *DLL4*. Conversely, HUVECs showed higher expression of genes linked to immune response and vascular homeostasis (e.g., *VCAM1*, and *ACE*) (Figure 4C; Supplementary Table 3).

**Figure 4:**
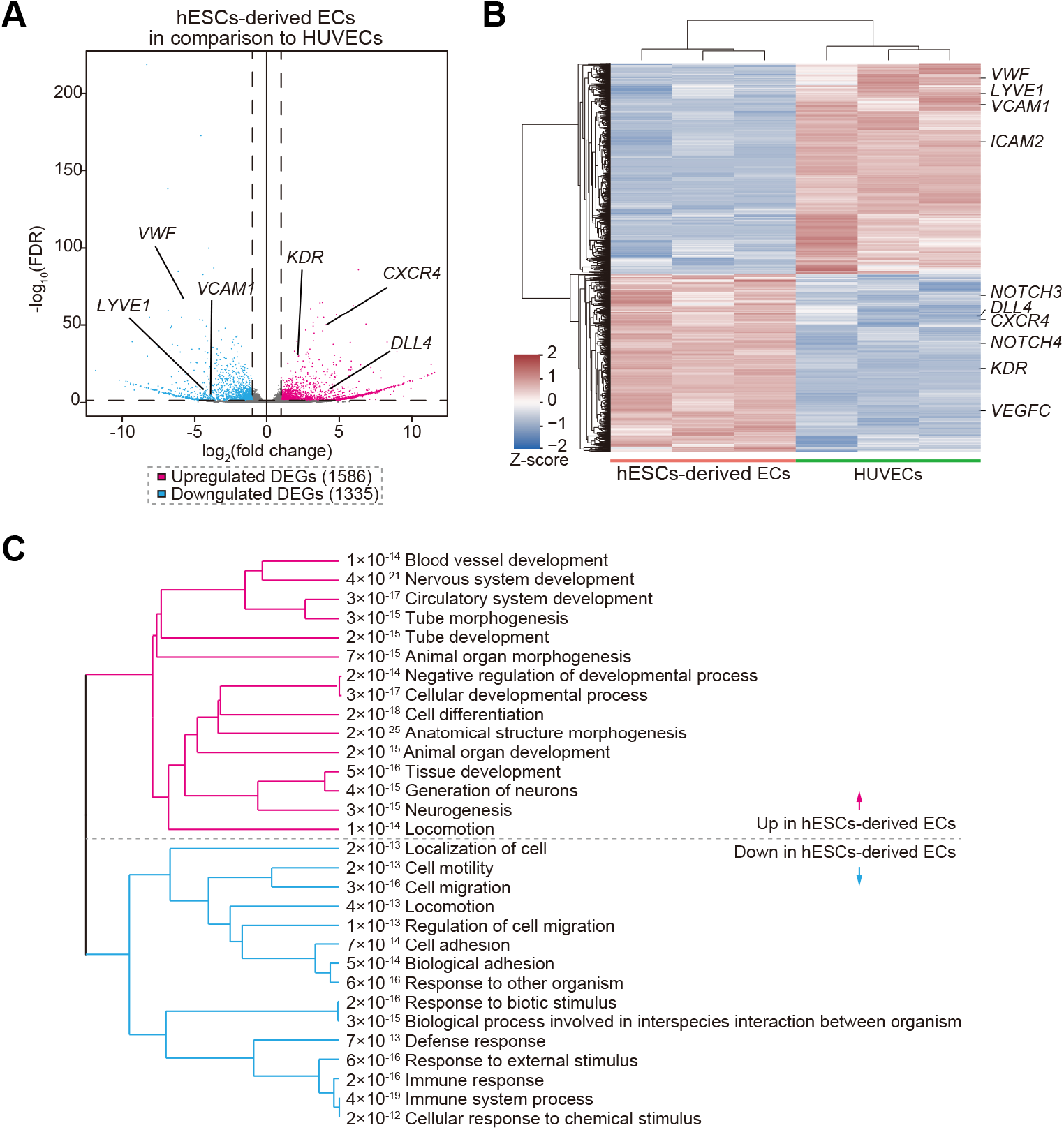
Differential gene-expression analysis of hESC-derived endothelial cells versus HUVECs (BRB-seq). (**A**) Volcano plot of differentially expressed genes (DEGs). Red points indicate genes up-regulated in hESC-derived ECs; blue points, genes up-regulated in HUVECs (|log_2_ fold-change| > 1, adj. *p* < 0.05). (**B**) Heatmap showing the expression of significant DEGs across individual hESC-EC and HUVEC samples. (**C**) Gene-ontology pathway enrichment of DEGs: pathways up-regulated in hESC-derived ECs (left) highlight vasculogenesis and tube morphogenesis, whereas those up-regulated in HUVECs (right) relate to immune response and external-stimulus signaling.

We confirmed that typical EC surface markers (*KDR, CD31, CD34*, and *CD144*) were similar between hPSC-derived ECs and HUVECs, but hPSC-derived ECs expressed lower levels of *TIE1, vWF*, and *NOS3* (Figure 5A). Moreover, the expression of arterial-specific genes (*NRP1, NOTCH1, CXCR4, DLL4*, and *EFNB2*) were significantly higher in hPSC-derived ECs, whereas venous genes (*NRP2*, and *EPHB4*) were lower (Figure 5B,C).

**Figure 5:**
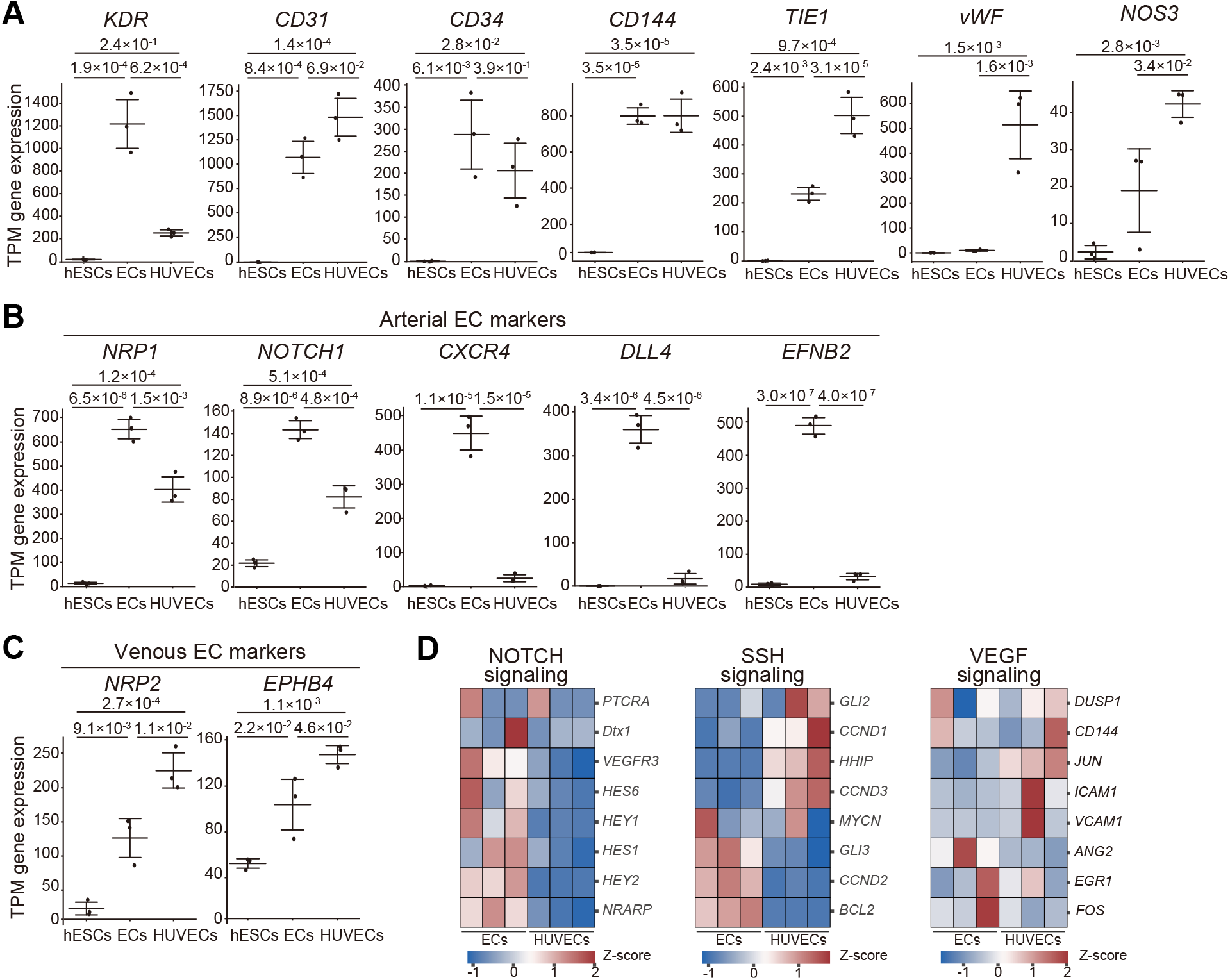
Endothelial marker and subtype-specific gene expression. (**A**) Transcripts per kilobase million (TPM) for core surface markers (*KDR, CD31, CD34, CD144*) and functional genes (*TIE1, vWF, NOS3*) in hESC-derived ECs, undifferentiated hESCs, and HUVECs. (**B**) TPM values for arterial markers (*NRP1, NOTCH1, DLL4, CXCR4, EFNB2*). (**C**) TPM values for venous markers (*NRP2, EPHB4*). Bars (or points) show mean ± s.d.; *P values were estimated* by Student’s *t*-test, and shown in the graphs. (**D**) Heatmap of NOTCH-, SHH-, and VEGF-signaling genes highlights stronger Notch-target activity in hESC-derived ECs, with no consistent differences in SHH or VEGF targets. Colors represent row-wise z-scores of TPM.

Pathway-level inspection showed stronger Notch-signaling activity in hPSC-derived ECs, whereas Shh and VEGF-target gene sets did not differ markedly (Figure 5D). Collectively, these data indicate that our protocol drives hPSCs toward an arterial-like endothelial phenotype, distinct from the venous identity of HUVECs.

## Discussion

In this study, we achieved highly efficient differentiation of hESCs into ECs with over 90% purity, using a protocol involving WNT pathway activation (CHIR99021) followed by NOTCH signaling inhibition (DAPT). Since *in vivo* studies have demonstrated that WNT signaling during mesoderm induction plays a critical role in EC differentiation,^23^ optimizing the duration of CHIR99021 treatment is necessary when differentiation is suboptimal. NOTCH signaling inhibitors promote differentiation into vascular endothelial progenitors by suppressing smooth muscle cell lineage commitment^33,34^. The resulting hESC-ECs display classic cobblestone morphology, efficiently take up acetylated LDL, and form capillary-like sprouts, providing clear evidence of endothelial identity. Bulk RNA barcoding and sequencing further revealed an arterial-skewed transcriptional profile: *NOTCH1, DLL4, CXCR4*, and *EFNB2* were elevated, whereas venous markers (*EPHB4*, and *NRP2*) were reduced. PSC-EC protocols often default to an arterial program unless venous cues are provided, suggesting that WNT activation coupled with endogenous VEGF signaling may override the transient NOTCH inhibition in our system. These results underscore that the combination of CHIR99021 and DAPT provides a simple, efficient route to generate endothelial cells, on par with or exceeding the efficiency of other state-of-the-art protocols.

The high yield and arterial bias of these ECs make them attractive for constructing vascularized organoids aimed at disease modeling and drug discovery, where perfusable microvessels improve organoid size, viability, and physiological relevance^11,28,35^. Because the cells can be generated in large numbers from pluripotent sources, including patient-specific iPSCs, the protocol also supports autologous cell therapies and the endothelialization of engineered grafts for ischemic disease or tissue repair^36^.

There are still limitations. First, the cells remain functionally immature, likely because they develop in static 2D culture lacking hemodynamic shear, extracellular matrix architecture, and crosstalk with mural or blood cells. Second, although enriched for arterial genes, the culture still contains a mixed endothelial population and offers no means to obtain venous ECs on demand. Finally, long-term stability, tumorigenic risk, and performance *in vivo* were not assessed and will require chemically defined, clinically compliant reagents.

To generate fully mature arterial or venous subtypes, stage-specific modulation of Notch and VEGF, exposure to laminar shear stress, or co-culture with pericytes should be tested. Post-differentiation conditioning, extended culture under flow, metabolic or epigenetic priming, or 3D matrix embedding, may boost vWF and NOS3 expression and enhance barrier and anti-thrombotic functions. Refining these parameters should yield endothelial cells that are not only abundant and phenotypically arterial or venous as needed but also functionally equivalent to their in vivo counterparts, accelerating the development of vascularized organoids, drug-screening platforms, and next-generation vascular therapies.

## Conclusions

We established the efficient method, brief WNT activation with CHIR99021 followed by Notch inhibition with DAPT, to differentiate hESCs into endothelial cells, resulting over 90 % purity without cell sorting. The cells show key endothelial behaviors (cobblestone shape, LDL uptake, sprouting) and an arterial-leaning gene profile. Although some mature markers remain lower than in primary HUVECs, the protocol yields large numbers of functional ECs suitable for vascularized organoids, drug testing, and future cell-therapy studies. Fine-tuning NOTCH/VEGF signaling and adding flow or co-culture steps should further mature the cells and allow selective arterial or venous production.

## List of abbreviations

ECs: Endothelial cells
hPSCs: human pluripotent stem cells
hESCs: human embryonic stem cells
hiPSCs: human induced pluripotent stem cells
HUVECs: human umbilical vein endothelial cells
FACS: fluorescence-activated cell sorting
MACS: magnetic-activated cell sorting
BRB-seq: bulk RNA barcoding and sequencing.

## Author Contributions

Conceptualization: KY, KK

Methodology: KY, ST, KK

Investigation: KY, KK

Visualization: KY

Funding acquisition: KY, KK

Project administration: KK

Supervision: KK

Writing – original draft: KY, KK

Writing – review & editing: KY, KK

## Availability of Data and Materials

All BRB-seq data are available at the Gene Expression Omnibus.

## Acknowledgments

We thank our lab members for their assistance. The authors thank to iCeMS Analysis Center to access the advanced microscopy and analytical instruments. We used ChatGPT o3 and Gemini 2.5 for correcting English grammar and readability.

