## Supplementary Figure 1 for "High-Purity Production of Endothelial Cells from Human Pluripotent Stem Cells"

**Koki Yoshimoto^1^, Shiho Terada^1^, Ken-ichiro Kamei**
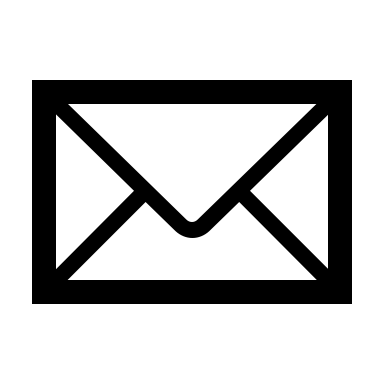
**,^1,2,3,4,5,6,7*^**

*^1^ Institute for Integrated Cell-Material Sciences, Kyoto University, Kyoto, 606-8501, Japan*

*^2^ Program of Biology, Division of Science, New York University Abu Dhabi, Abu Dhabi, P.O. Box 129188, United Arab Emirates*

*^3^ Program of Bioengineering, Division of Engineering, New York University Abu Dhabi, Abu Dhabi, P.O. Box 129188, United Arab Emirates*

*^4^ Department of Biomedical Engineering, Tandon School of Engineering, New York University, New York, NY 10003, USA*

*^5^ Department of Biology, Faculty of Arts & Science, New York University, Brooklyn, NY 11201, USA*

*^6^ Department of Pharmaceutics, Wuya College of Innovation, Shenyang Pharmaceutical University, Shenyang, 110016, China*

*^7^ Joint International Research Laboratory of Intelligent Drug Delivery Systems, Ministry of Education, Shenyang, 110016, China*

Address correspondence to: Ken-ichiro Kamei, PhD

Address: New York University Abu Dhabi (NYUAD), Saadiyat Campus, #C1-032, P.O. Box 129188, Abu Dhabi, United Arab Emirates

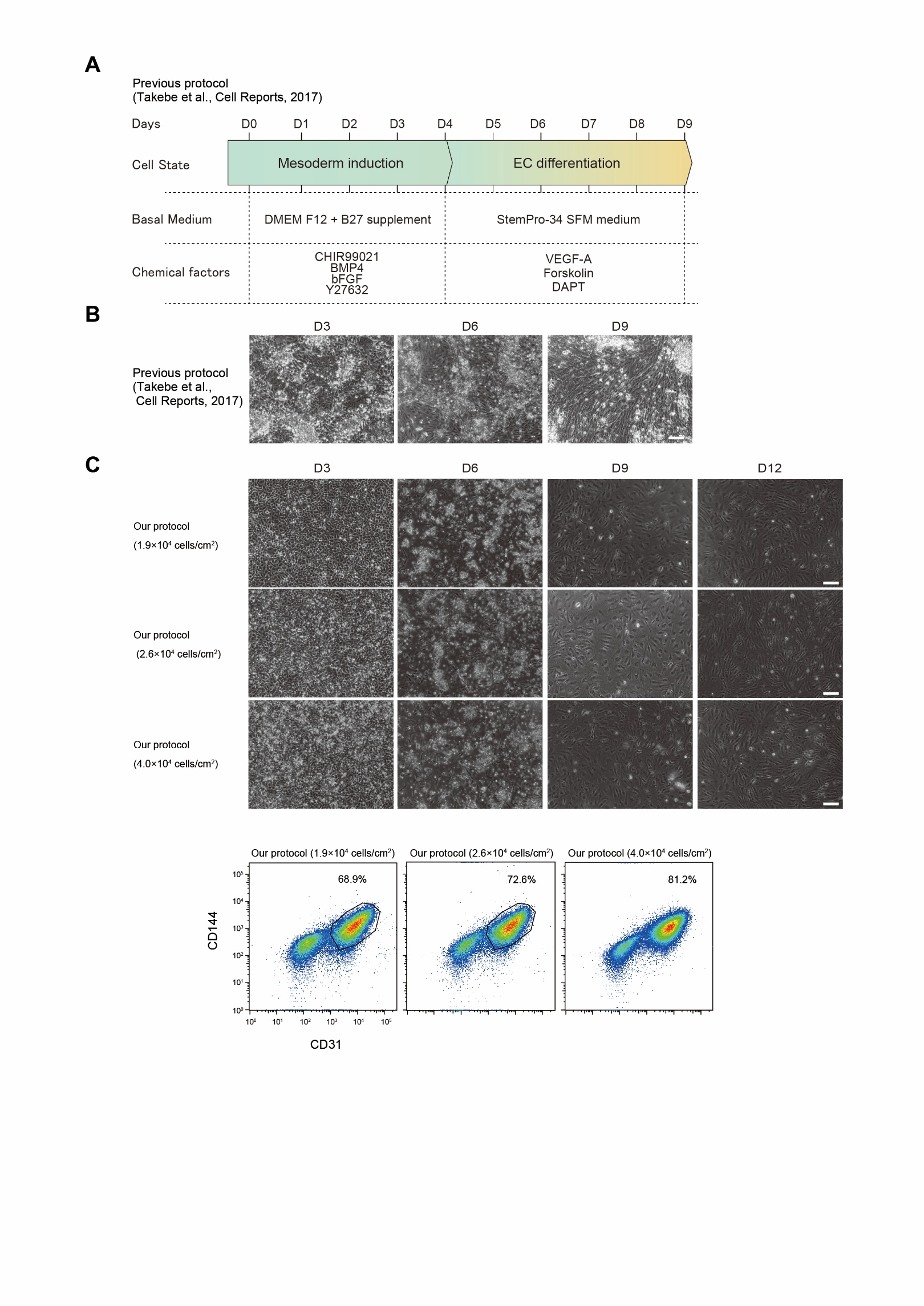

**Supplementary Figure 1:** Comparison of endothelial differentiation protocols. (**A**) Endothelial differentiation protocols previously reported by Takebe et al. (**B**) Representative photographs of the obtained cells with previously reported protocol. Scale bar represents 100 µm. (**C**) Representative photographs of the obtained cells with our protocol on the three density conditions. Scale bars represent 100 µm.
